# Glutamine-Dependent Downregulation of FLT3-ITD is a Mechanism of FLT3 Inhibitor Resistance in FLT3-ITD AML in Hypoxia

**DOI:** 10.64898/2026.05.02.722336

**Authors:** Giovannino Silvestri, Aditi Chatterjee, Blair P. Rendina, Eli E. Bar, Maria R. Baer

## Abstract

FLT3 inhibitors have improved outcomes in acute myeloid leukemia (AML) with *FMS*-like tyrosine kinase 3 internal tandem duplication (FLT3-ITD), but responses are not durable. Notably, FLT3 inhibitors clear blasts from the blood, but not the bone marrow, a hypoxic niche. We investigated effects of hypoxia and the key nutrient glutamine on FLT3 inhibitor response. FLT3-ITD AML cell lines and patient blasts were cultured with FLT3 inhibitors under normoxia (21%) or hypoxia (<1% O_2_) with or without glutamine or the glutaminase inhibitor telaglenastat (CB-839). Cytotoxicity was measured in WST-1 assays and drug combination effects by Chou-Talalay analysis. Protein expression was measured by immunoblotting, turnover and proteasomal degradation by cycloheximide chase with and without MG-132, and mRNA expression by RT-qPCR. Effect of the ubiquitin ligase c-CBL was tested by siRNA knockdown. FLT3 inhibitor IC_50_s were 3-5-fold higher in hypoxia than normoxia, associated with FLT3-ITD and p-STAT5 downregulation and accelerated FLT3-ITD proteasomal degradation (half-life, 1.0 vs. 2.5 hours). c-CBL expression increased in hypoxia, and c-CBL knockdown restored FLT3-ITD expression and FLT3 inhibitor sensitivity. Glutamine deprivation or telaglenastat treatment abrogated c-CBL upregulation in hypoxia and preserved FLT3-ITD and p-STAT5 expression and FLT3 inhibitor sensitivity. Telaglenastat synergized with FLT3 inhibitors in hypoxia, supporting clinical testing.

**Graphical Abstract:** 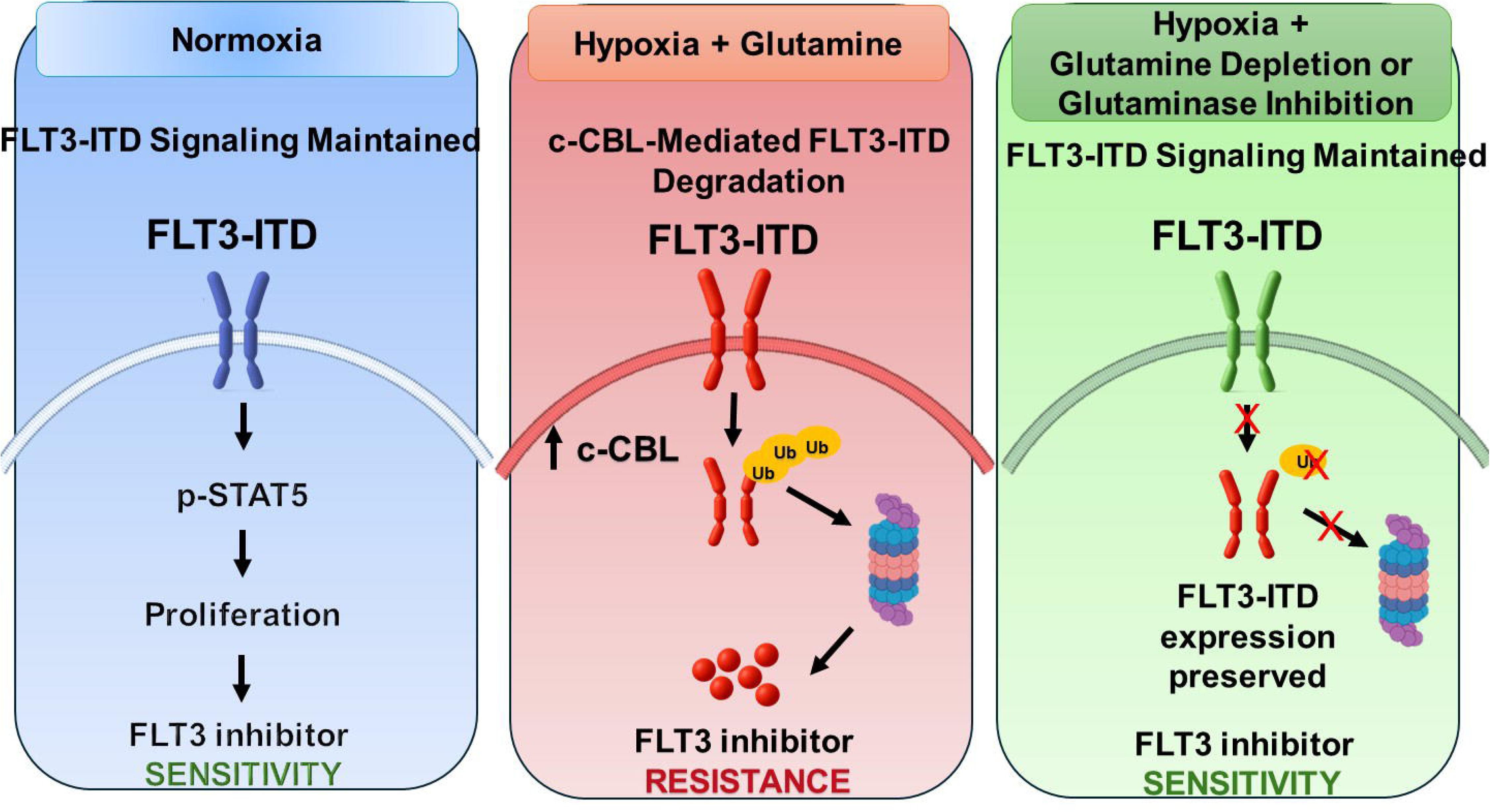

## Introduction

Internal tandem duplication in the *FMS*-like tyrosine kinase 3 gene (*FLT3*-ITD) is present in blasts of approximately 25% of patients with acute myeloid leukemia (AML), historically associated with poor patient outcomes (1–3). FLT3-ITD constitutively activates the FLT3 receptor tyrosine kinase (4). Wild-type (WT) FLT3 signals through PI3 kinase (PI3K)-AKT-mTOR and Ras-Raf-MEK-ERK upon binding of FLT3 ligand (4). FLT3-ITD signals constitutively through these pathways, and also, aberrantly, though signal transducer and activator of transcription (STAT) 5, promoting AML cell survival, proliferation and resistance to apoptosis (4).

FLT3-ITD AML patient outcomes have improved with incorporation of FLT3 tyrosine kinase inhibitors into treatment (5), but remain suboptimal. Treatment with the FLT3 inhibitors midostaurin or quizartinib in conjunction with chemotherapy improved disease-free (6), relapse-free (7) and overall (6,7) survival, compared to placebo, in FLT3-ITD AML patients, but long-term survival rate was only 50% (6,7). Similarly, gilteritinib improved remission rate and survival in patients with relapsed/refractory FLT3-ITD AML, compared to chemotherapy, but the complete remission rate with gilteritinib was only 20.5%, and median survival only 9.3 months (8).

Notably, FLT3 inhibitor treatment clears FLT3-ITD AML blasts from the peripheral blood (PB), but not from the bone marrow (BM) (9), an apparent protective niche that impairs response to FLT3 inhibitors. Therefore treatment strategies to eradicate FLT3-ITD AML must not only target AML cells, but also disrupt protective cues from the BM niche, where FLT3-ITD AML cells persist and may then also acquire mutations conferring intrinsic FLT3 inhibitor resistance (10).

The BM niche is a complex network comprised of mesenchymal stromal cells, endothelial cells, osteoblasts, immune cells and extracellular matrix components, which collectively create a supportive environment through both direct cell-cell interactions and secretion of soluble factors (11,12). A defining feature of the niche is physiologic hypoxia, with oxygen tensions as low as 1% in endosteal and perivascular regions (13). Regional hypoxic zones harbor hematopoietic stem cells (14), as well as quiescent AML cells which resist therapy and persist as residual disease (15).

Nutrient composition also impacts adaptation of leukemia cells to the BM niche. Glutamine promotes oxidative phosphorylation in AML cells, and supports proliferation and resistance to apoptosis (16). FLT3-ITD AML cells, in particular, are metabolically dependent on glutamine, and deletion of glutaminase, the first enzyme in glutamine metabolism, was found to be synthetically lethal with FLT3 inhibition (17). Moreover, in another study, glutamine was implicated in FLT3 inhibitor resistance, with l-asparaginase, which hydrolyzes glutamine, as well as the glutaminase inhibitor CB-839, sensitizing FLT3-ITD AML cells to quizartinib (18).

We investigated the roles of hypoxia and of glutamine availability in FLT3 inhibitor resistance, as an approach to understanding and overcoming FLT3 inhibitor resistance of FLT3-ITD AML cells in the BM niche.

## Materials and methods

### Cell lines

The human FLT3-ITD AML cell lines MV4-11 and MOLM-14, with homozygous and heterozygous FLT3-ITD, respectively (American Type Culture Collection, Manassas, VA, USA) were maintained in RPMI 1640 medium (Gibco, Grand Island, NY, USA) with 10% fetal bovine serum, 1% penicillin-streptomycin and 2 mM L-glutamine, unless otherwise indicated, and were tested for Mycoplasma every six months (19).

### Patient samples

Blood samples were obtained from patients with FLT3-ITD AML (Supplementary Table S1) on a University of Maryland School of Medicine Institutional Review Board-approved protocol, following written informed consent. Studies were conducted in accordance with the Declaration of Helsinki. Blasts were isolated by density centrifugation over Lymphocyte Separation Medium (Corning, Glendale, AZ, USA, C-44010) and cultured in StemSpan CC100 medium (STEMCELL Technologies, Vancouver, BC, CAN, 02690) supplemented with cytokines [20 ng/mL interleukin (IL)-3, 20 ng/mL IL-6, 100 ng/mL stem cell factor, 100 ng/mL FLT3 ligand] and 2 mM L-glutamine, unless otherwise indicated.

### Materials

The FLT3 inhibitors gilteritinib (ASP 2215; S7754) (Type I) and quizartinib (AC220, S1526) (Type II) were from Selleck Chemicals, Houston, TX, USA, and the glutaminase inhibitor telaglenastat (CB-839; MedChemExpress, Monmouth Junction, NJ, USA, HY-12248).

### Cell cultures

FLT3-ITD AML cell lines and primary blasts were cultured in normoxia (21% O_2_) or hypoxia (<1% O_2_; 94% N_2_), with 5% CO_2_, at 37°C for up to 96 hours using the Xvivo System Model X3 (BioSpherix, Redfield, NY, USA). For time course studies, cells were harvested at 0, 48 and 96 hours. For glutamine modulation studies, cells were cultured with or without 2 mM L-glutamine or with 2 mM L-glutamine and 100 nM telaglenastat.

### Cytotoxicity assay

Cytotoxicity was measured using the WST-1 assay (19). Cells were plated in triplicate at 20,000/well in 96-well plates and treated with gilteritinib or quizartinib at serial concentrations for 48 or 96 hours in normoxia or hypoxia, with or without 2 mM glutamine. WST-1 reagent (Millipore Sigma, Burlington, MA, USA, 11644807001) was added and cells were incubated for 2-3 hours at 37°C. Absorbance at 450 nm was measured using a microplate reader. IC_50_ values were calculated by non-linear regression analysis using Prism 11 software (GraphPad, San Diego, CA, USA).

### Immunoblotting

Cells harvested at serial time points were lysed in RIPA buffer (150 mM NaCl) with protease/phosphatase inhibitors (Roche, Indianapolis, IN, USA, 11836170001). Protein concentration was determined using the Quick Start Bradford protein assay kit (Bio-Rad, Hercules, CA, USA, 5000201). Protein (25 µg for cell lines, 35 µg for patient samples) was separated by SDS-PAGE, transferred to PVDF membranes, and probed with antibodies to FLT3 (3462S), p-STAT5 (Y694) (9351), STAT5 (94205), c-CBL (2747), p-c-CBL (Y371) (3554) (Cell Signaling Technology, Danvers, MA, USA) and vinculin (Millipore Sigma, V9264) and horseradish peroxidase-conjugated secondary antibodies. Signals were detected by enhanced chemiluminescence (Thermo Fisher Scientific, Waltham, MA, USA, 34094, 21106, 34578). Band intensities were quantified by densitometry (Image J, NIH) and normalized to vinculin and, for time course studies, to Time 0 levels.

### Protein turnover and proteasomal degradation

Cells cultured in normoxia or hypoxia were treated with cycloheximide (CHX, 100 µg/mL; Millipore Sigma, C7698) for 60 minutes to inhibit protein synthesis and harvested at 0, 1, 2, and 4 hours, starting one hour after addition of CHX. For proteasomal degradation studies, cells were treated with CHX with or without 20 µM proteasome inhibitor carbobenzoxy-L-leucyl-L-leucyl-L-leucinal (MG-132; Millipore Sigma, 474790) added 30 minutes after CHX. Protein expression was analyzed by immunoblotting. Band intensities quantified by densitometry were normalized to vinculin and Time 0. Protein half-lives were determined by linear regression using Prism 11 software (GraphPad) (20).

### Quantitative real-time polymerase chain reaction (RT-qPCR)

Total RNA was extracted using RNeasy Plus Mini Kits (Qiagen, Germantown, MD, USA, 74134) and quantified using a NanoDrop™ Lite Spectrophotometer (Thermo Fisher Scientific). cDNA was synthesized from 0.5-1 µg RNA using iScript (Bio-Rad, 1708896). qRT-PCR was performed with SsoAdvanced Universal SYBR Green Supermix (Bio-Rad, 725271EDU) on a CFX96 (Bio-Rad) (95 °C for 3 minutes, 40 cycles of 95°C for 10 seconds, 60°C for 30 seconds; melting curve). Relative expression was calculated by ΔΔCt, with glyceraldehyde 3-phosphate dehydrogenase (GAPDH) RNA as reference. FLT3, c-CBL and GAPDH primers are listed in Supplementary Table S2.

### c-CBL silencing

Transient knockdown of the E3 ubiquitin ligase Casitas B-lineage Lymphoma (c-CBL) was performed using siGENOME SMARTpool small interfering RNA (siRNA) (Horizon Discovery, Cambridge, UK, M-003003-02-0050). Cells were seeded at 30,000/well in 24-well plates containing 500 µL antibiotic-free complete medium. Plates were incubated overnight at 37°C in a humidified 5% CO_2_ incubator. Cells were transfected with c-CBL or control siRNA using DharmaFECT following manufacturer’s protocol and analyzed 48-96 hours post-transfection. qPCR and immunoblotting were performed to verify knockdown efficiency.

### Drug combination studies

Cells seeded in 96-well plates were treated in triplicate with gilteritinib or quizartinib and telaglenastat at diverse concentrations alone and in combinations, and cytotoxicity was measured using the WST-1 assay, as above (19). Drug combination effects were analyzed by the Chou-Talalay method, using CompuSyn software (Paramus, NJ, USA) (19). Combination index values <1, =1 or >1 indicate synergy, additivity, or antagonism, respectively (21).

### Statistical analysis

All experiments were performed in at least two independent biological replicates unless otherwise specified. Data are presented as mean ± standard error of the mean. Statistical analyses were conducted using unpaired two-tailed Student’s t-tests or one-way analysis of variance with appropriate post hoc tests. p<0.05 was considered significant.

## Results

### FLT3-ITD AML cells are resistant to FLT3 inhibitors in hypoxia

To determine the impact of hypoxia on FLT3 inhibitor sensitivity, we cultured primary FLT3-ITD AML blasts from two patients with gilteritinib or quizartinib in increasing concentrations for 96 hours in normoxia (21% O_2_) or hypoxia (<1% O_2_). Cytotoxicity of both gilteritinib and quizartinib was reduced 2-3-fold in hypoxia, compared to normoxia (Figure 1A).

**Figure 1.**
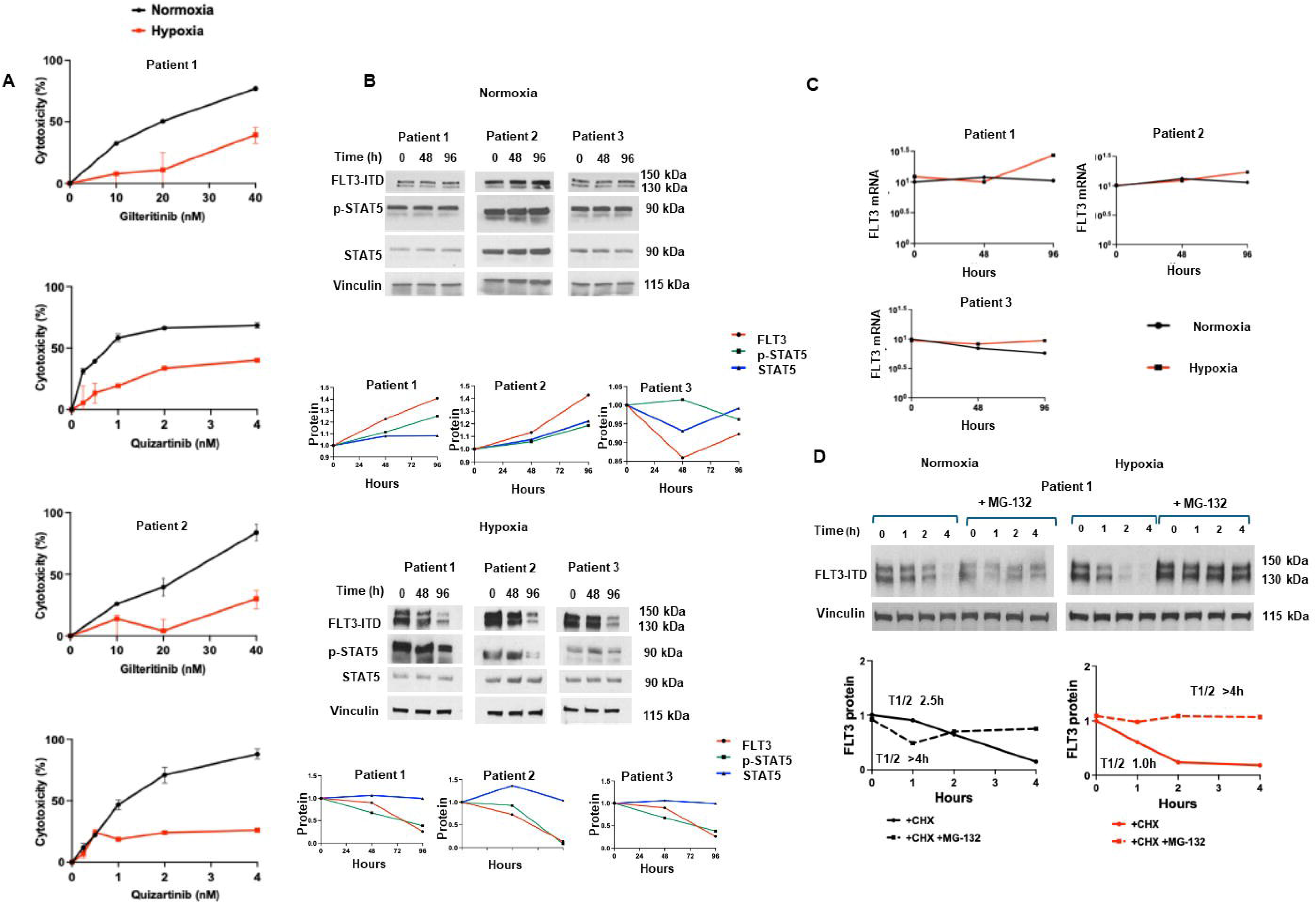
FLT3-ITD AML cells are resistant to FLT3 inhibitors in hypoxia, associated with post-translational FLT3-ITD downregulation. A. Primary FLT3-ITD AML blasts from two patients were cultured for 48 hours under normoxia (21%O_2_) or hypoxia (<1% O_2_) with the FLT3 inhibitors gilteritinib or quizartinib at increasing concentrations and cytotoxicity was measured using the WST-1 assay. Cytotoxicity of both gilteritinib and quizartinib was reduced in hypoxia (red lines) compared with normoxia (black lines). Data represent means of three replicate wells. B. Blasts from three FLT3-ITD AML patients cultured in normoxia and hypoxia were harvested at 0, 48 and 96 hours and immunoblotted for expression of FLT3, p-STAT5, STAT5 and vinculin loading control. Bands were quantified by densitometry and FLT3, p-STAT5 and STAT5 were normalized to vinculin and to Time 0. Immunoblots and graphs are shown. FLT3-ITD and p-STAT5 were downregulated in hypoxia, but not in normoxia, while total STAT5 remained stable in both. C. FLT3 and GAPDH control mRNA was measured by RT-qPCR in blasts from three FLT3-ITD AML patients cultured for 0, 48 and 96 hours in hypoxia and normoxia. Graphs of FLT3 mRNA, normalized to GAPDH mRNA, show no decrease in FLT3 mRNA. D. FLT3-ITD AML patient blasts were treated with cycloheximide (CHX, 100 µg/mL) to block new protein translation, with or without addition of the proteasome inhibitor MG-132 (20 µmol/L) after 30 minutes to block proteasomal degradation, then harvested at 0, 1, 2, and 4 hours, starting 1 hour after addition CHX, and immunoblotted for FLT3 and vinculin control. Bands were quantified by densitometry and FLT3 protein half-lives were calculated by linear regression analysis. FLT3-ITD protein turnover was accelerated in hypoxia (1.0 vs. 2.5 hours), in the absence, but not presence, of MG-132, consistent with increased proteasomal degradation in hypoxia.

### FLT3-ITD expression and STAT5 activation are down-modulated in hypoxia

Expression of FLT3-ITD and downstream p-STAT5 (Y694), as well as total STAT5, was measured by immunoblotting in primary AML blasts from three FLT3-ITD AML patients cultured in normoxia or hypoxia for 96 hours. FLT3-ITD, p-STAT-5, and STAT5 expression did not change in 96-hour culture in normoxia (Figure 1B). In contrast, FLT3-ITD and p-STAT5 expression decreased progressively over time in all three patient samples in hypoxia, while total STAT5 remained unchanged, confirmed by densitometric analyses (Figure 1B).

### FLT3-ITD downregulation in hypoxia is not transcriptional

We next examined whether downregulation of FLT3-ITD expression in hypoxia is transcriptional. FLT3 mRNA levels were unchanged over 96 hours in hypoxia, indicating a non-transcriptional mechanism (Figure 1C).

### FLT3-ITD proteasomal degradation is accelerated in hypoxia

To test whether hypoxia-induced FLT3-ITD downregulation is post-translational, CHX chase assays were performed in primary FLT3-ITD AML blasts cultured in normoxia and hypoxia. CHX (100 µg/mL) was added to cell cultures to block new protein translation, and cells were harvested for immunoblotting at 0, 1, 2, and 4 hours, starting 1 hour after addition CHX. FLT3-ITD protein half-life was shorter in cells cultured in hypoxia, compared to normoxia (1.0 vs. 2.5 hours) (Figure 1D). Cells cultured with the proteasome inhibitor MG-132 added 30 minutes after addition of CHX showed abrogation of FLT3-ITD turnover in hypoxia and normoxia (half-lives >4 hours in both) (Figure 1D), demonstrating that FLT3-ITD turnover in hypoxia, as well as normoxia, is proteasome-dependent.

### c-CBL expression is increased in hypoxia

The E3 ubiquitin ligase c-CBL is a key regulator of turnover of receptor tyrosine kinases, including FLT3, promoting their ubiquitination and proteasomal degradation (22). c-CBL expression has been reported to increase in cardiomyocytes in hypoxia (23), suggesting that it might also increase in FLT3-ITD AML blasts in hypoxia and promote FLT3-ITD proteasomal degradation.

Total and phosphorylated (Y371) c-CBL (24) expression was measured in blasts from three FLT3-ITD AML patients cultured in normoxia or hypoxia for 96 hours. A consistent time-dependent increase in both total c-CBL and p-c-CBL was observed in all three patient samples in hypoxia, reaching approximately 2.5-3.5-fold by densitometric analysis, whereas no sustained increase was observed in normoxia (Figure 2A). These results suggested that c-CBL could play a central role in mediating increased FLT3-ITD proteasomal degradation in hypoxia.

**Figure 2.**
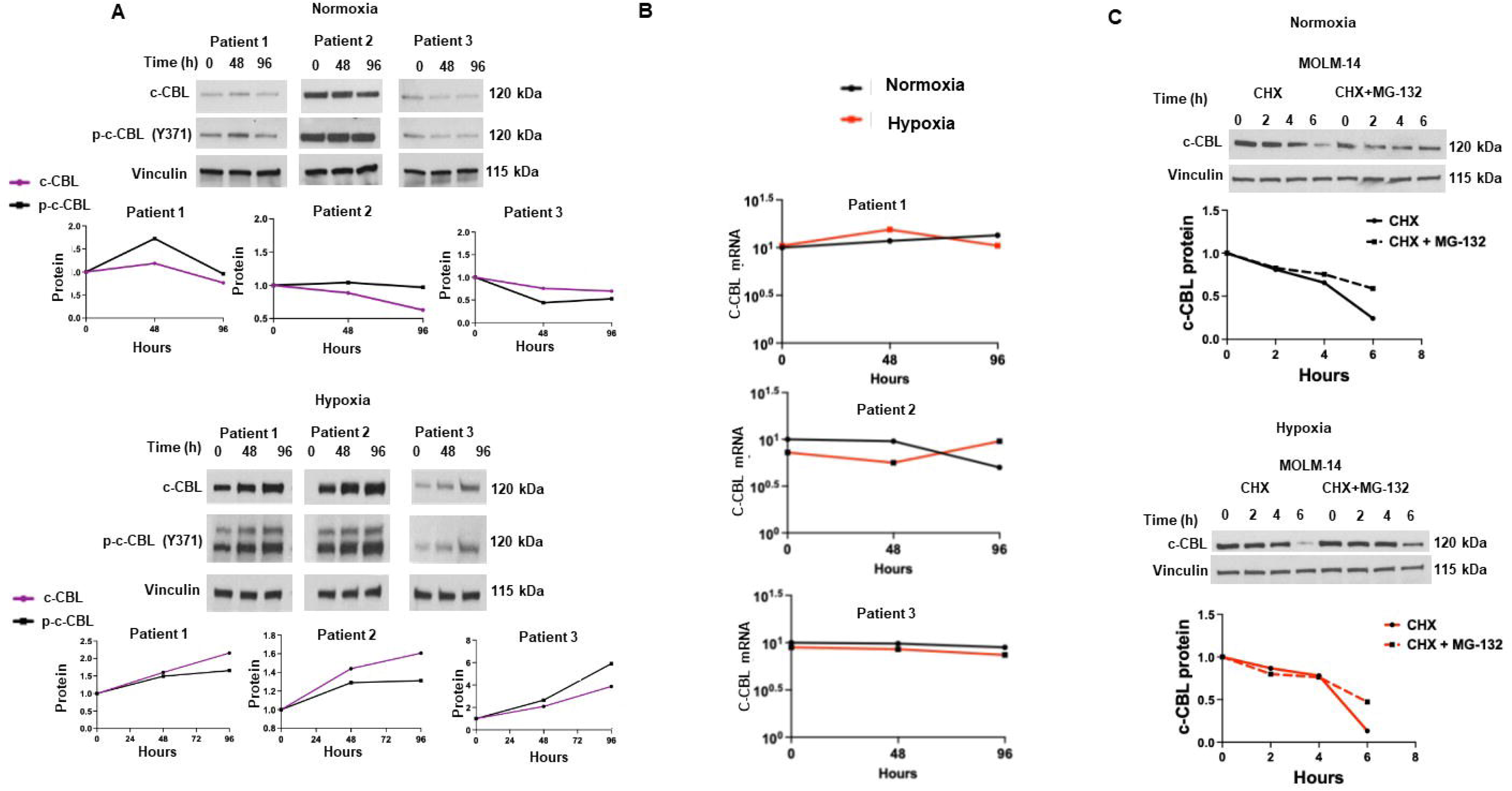
Expression of the ubiquitin ligase c-CBL is increased in hypoxia. **A.** Blasts from three FLT3-ITD AML patients cultured in normoxia and hypoxia were harvested at 0, 48 and 96 hours and immunoblotted for expression of c-CBL, p-c-CBL (Y371) and vinculin loading control, and bands were quantified by densitometry. Expression of c-CBL and p-c-CBL was upregulated in hypoxia, but not in normoxia. **B.** c-CBL and GAPDH control mRNA was measured at 0, 48 and 96 hours by RT-qPCR, and graphs of c-CBL mRNA normalized to GAPDH control mRNA are shown. c-CBL mRNA expression did not decrease in cells cultured in hypoxia. **C**. MOLM-14 cells were treated with cycloheximide (CHX, 100 µg/mL), with or without addition of the proteasome inhibitor MG-132 (20 µmol/L) after 30 minutes, then harvested at 0, 1, 2, and 4 hours, starting 1 hour after addition CHX, and immunoblotted for FLT3 and vinculin control. Bands were quantified by densitometry and FLT3 protein half-lives were calculated by linear regression analysis. FLT3-ITD protein turnover was similar in hypoxia and normoxia and the effect of MG-132 was also similar.

To determine whether the increase in c-CBL expression in hypoxia was transcriptionally regulated, c-CBL mRNA expression was measured by RT-qPCR in cells cultured in normoxia or hypoxia for 96 hours. c-CBL mRNA levels did not increase over time in hypoxia and were similar to those observed in normoxia (Figure 2B), indicating that c-CBL upregulation is not transcriptionally mediated.

We next assessed whether altered protein stability contributed to increased c-CBL levels in hypoxia. Cycloheximide chase experiments performed in normoxia and hypoxia showed similar c-CBL degradation kinetics, and similar effects of the proteasome inhibitor MG-132 (Figure 2C). Based on these results, the increase in c-CBL protein levels in hypoxia is not due to altered protein stability or proteasomal degradation.

### c-CBL knockdown abrogates FLT3-ITD downregulation in hypoxia

To test the functional role of c-CBL in FLT3-ITD downregulation in hypoxia, siRNA-mediated silencing of c-CBL was performed in the MOLM-14 and MV4-11 FLT3-ITD AML cell lines and in one primary FLT3-ITD AML patient sample (Figure 3A). MOLM-14, MV4-11 and primary FLT3-ITD AML patient cells transfected with c-CBL siRNA or control siRNA were cultured in hypoxia for 96 hours, harvested at 0, 48 and 96 hours and immunoblotted for expression of c-CBL, FLT3-ITD and vinculin loading control. Time-dependent decrease in FLT3-ITD protein expression was seen in cells treated with siRNA control, while c-CBL knockdown abrogated this time-dependent FLT3-ITD downregulation, maintaining stable expression at 48 and 96 hours (Figure 3B). c-CBL mRNA expression decreased in MOLM-14, MV4-11 and primary FLT3-ITD AML patient cells transfected with c-CBL siRNA, but not control siRNA, as expected, cultured for 96 hours, but there was no effect on FLT3 mRNA expression (Figure 3C). FLT3-ITD protein turnover and proteasomal degradation were then studied in MOLM-14 cells transfected with c-CBL and control siRNA. FLT3-ITD protein proteasomal degradation in hypoxia was abrogated by c-CBL silencing (Figure 3D). The data are consistent c-CBL mediating increased FLT3 proteasomal degradation as a mechanism of FLT3-ITD downregulation in hypoxia.

**Figure 3.**
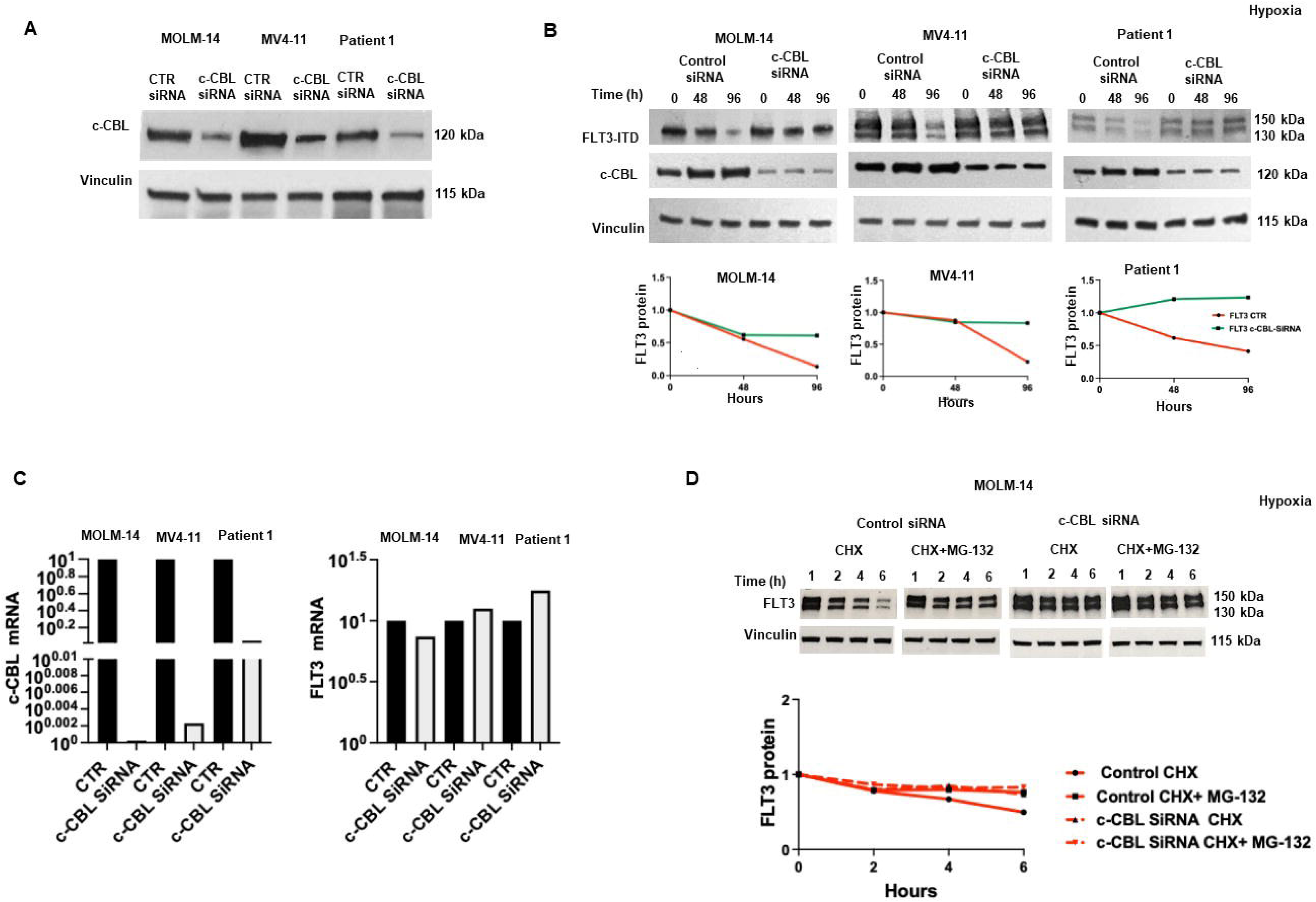
c-CBL knockdown abrogates FLT3-ITD downregulation in hypoxia. **A.** Whole-cell lysates of MOLM-14 and MV4-11 FLT3-ITD AML cells and primary FLT3-ITD AML blasts (Patient 1) transfected with c-CBL siRNA or control (CTR) siRNA were immunoblotted and probed for c-CBL and vinculin loading control, and bands were quantified by densitometry. Immunoblots demonstrate c-CBL knockdown in cells transfected with c-CBL siRNA. **B.** MOLM-14 and MV4-11 FLT3-ITD AML cell lines and primary FLT3-ITD AML patient blasts transfected with c-CBL siRNA or CTR siRNA cultured in hypoxia were harvested at 0, 48, or 96 hours, and whole-cell lysates were probed for FLT3, c-CBL and vinculin loading control. Immunoblots and densitometric analysis are shown, demonstrating that c-CBL knockdown abrogates FLT3-ITD downregulation in hypoxia. **C.** c-CBL, FLT3 and GAPDH control mRNA expression was measured by RT-qPCR in MOLM-14, MV4-11 and primary FLT3-ITD AML patient blasts treated with c-CBL siRNA or CTR siRNA for 96 hours. Data are shown graphically, with bars showing mean fold expression (2^-ΔCt) ± SD in cells transfected with c-CBL siRNA, relative to CTR siRNA. c-CBL silencing decreases c-CBL, but not FLT3, mRNA, showing that, as expected, FLT3 downregulation in cells transfected with CTR siRNA is not transcriptional. **D.** MOLM-14 cells transfected with c-CBL siRNA or CTR siRNA were treated with CHX to inhibit new protein translation, with or without MG-132 to inhibit proteasomal degradation. FLT3 protein turnover was accelerated in cells transfected with control, but not c-CBL, siRNA, in the absence, but not presence, of MG-132, consistent with c-CBL silencing rescuing FLT3-ITD AML cells from increased FLT3-ITD proteasomal degradation in hypoxia.

### Modulation of FLT3 expression is heterogeneous in WT FLT3 AML blasts in hypoxia

To test whether downregulation of FLT3 expression in hypoxia is specific for cells with FLT3-ITD, blasts from three patients with AML with WT FLT3 cultured in hypoxia for 96 hours were immunoblotted for FLT3, c-CBL and vinculin loading control (Figure 4). A slight progressive decrease in FLT3 protein levels was observed in blasts from Patient 7 over time, whereas FLT3 expression remained relatively stable in blasts from Patients 8 and 9, indicating interpatient heterogeneity in hypoxia-induced receptor regulation. c-CBL protein levels also showed modest variation across samples without a consistent trend. These findings suggest that hypoxia does not uniformly induce FLT3 downregulation nor c-CBL upregulation in AML blasts with WT FLT3, in contrast to FLT3-ITD.

**Figure 4.**
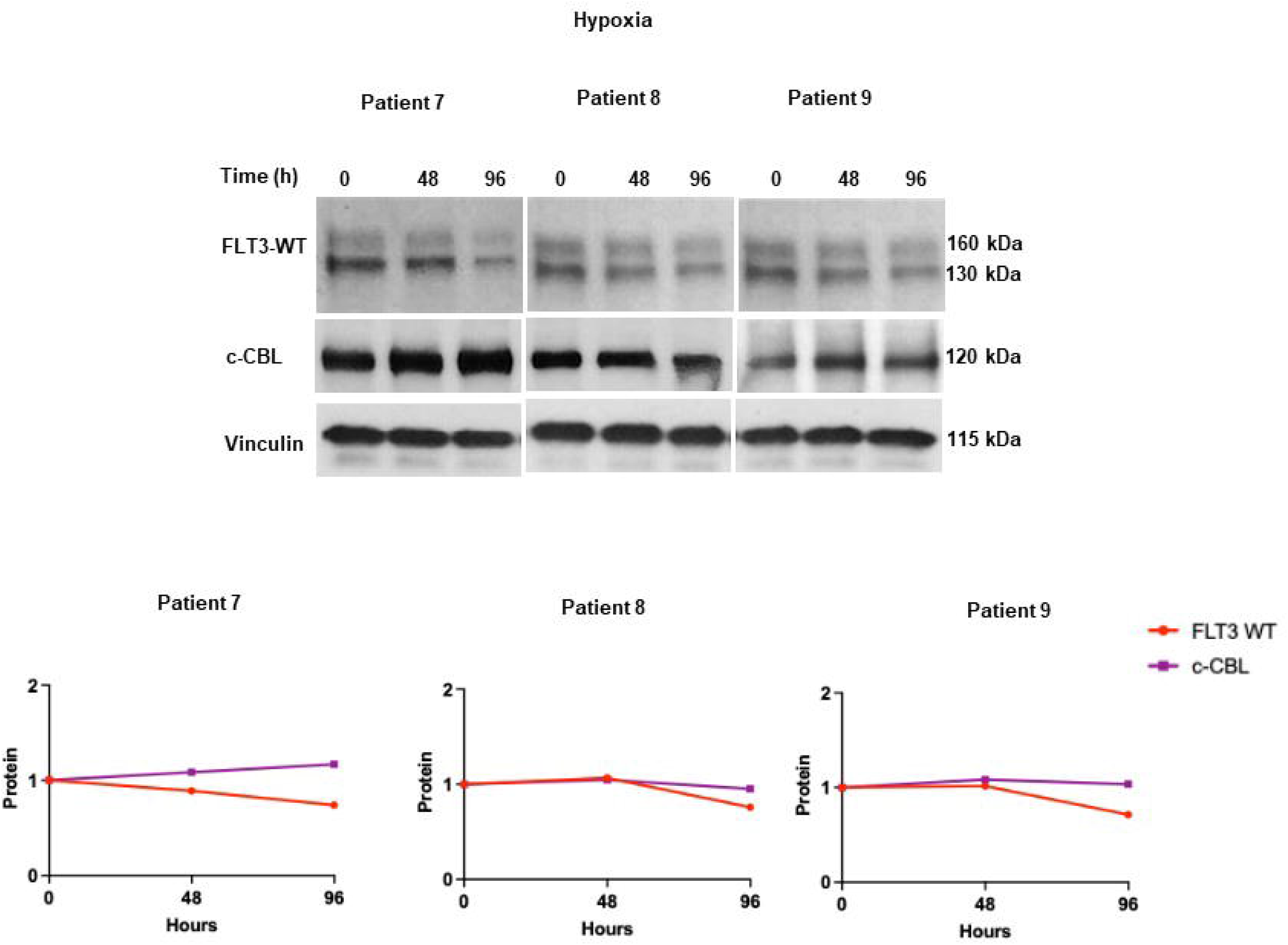
Primary AML blasts with WT FLT3 exhibit heterogeneous modulation of FLT3 expression in hypoxia. Primary blasts from three patients with AML with WT FLT3 (Patients 7–9) cultured in hypoxia for 96 hours were immnunoblotted for FLT3, c-CBL and vinculin loading control. The findings suggest that, in contrast to FLT3-ITD AML, hypoxia does not uniformly induce FLT3 downregulation in FLT3-WT primary blasts, and that additional patient-specific factors may influence FLT3 stability under low-oxygen conditions.

### FLT3-ITD AML cells remain sensitive to FLT3 inhibitors in hypoxia in the absence of glutamine, associated with preserved FLT3-ITD and p-STAT5 expression

Because the BM microenvironment is glutamine-rich and glutamine has been shown to impact FLT3 inhibitor sensitivity (17,18), we investigated the effect of glutamine on FLT3 inhibitor sensitivity and FLT3-ITD expression in hypoxia.

FLT3-ITD AML blasts from four patients were cultured in medium with gilteritinib or quizartinib at escalating concentrations with or without 2 mM L-glutamine, in normoxia or hypoxia for 96 hours. Cytotoxicity of both gilteritinib and quizartinib was decreased in hypoxia, but not normoxia, in the presence of glutamine, while cytotoxicity of both drugs in cells cultured in hypoxia in the absence of glutamine was similar to that of cells cultured in normoxia (Figure 5A).

**Figure 5.**
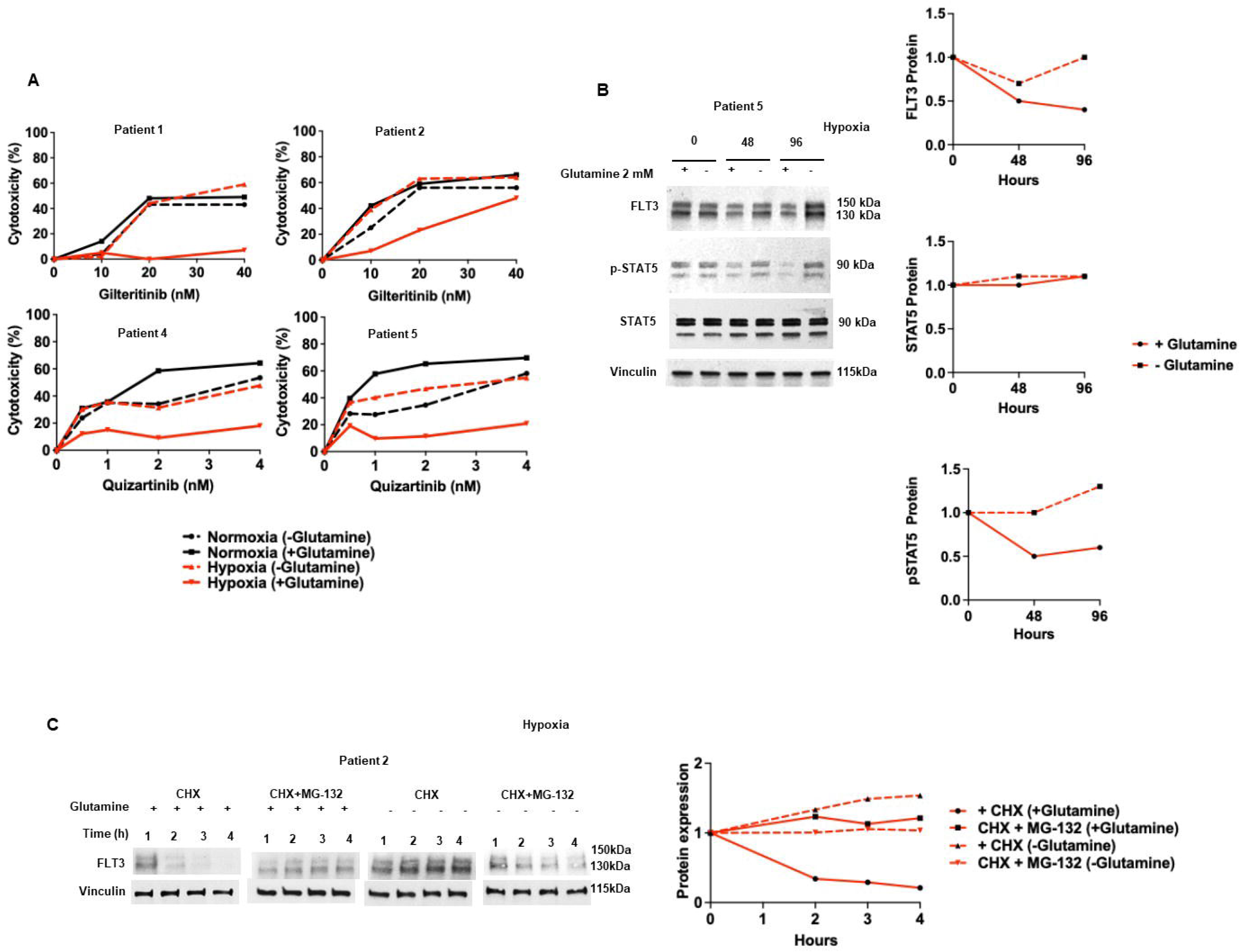
FLT3-ITD cells remain sensitive to FLT3 inhibitors in hypoxia in the absence of glutamine, associated with preserved FLT3-ITD and p-STAT5 expression. **A.** Primary FLT3-ITD AML cells from four patients were cultured for 48 hours with the FLT3 inhibitors gilteritinib or quizartinib in increasing concentrations in normoxia or hypoxia in the presence or absence of 2 mM glutamine, and cytotoxicity was measured using the WST-1 assay. FLT3-ITD AML cells remained sensitive to FLT3 inhibitors in hypoxia in the absence of glutamine. **B.** Primary FLT3-ITD AML cells cultured in hypoxia in the presence or absence of 2 mM glutamine were harvested for immunoblotting at 0, 48 and 96 hours. FLT3-ITD and p-STAT5 expression decreased progressively in the presence, but not absence, of 2 mM glutamine, while total STAT5 expression did not change, demonstrating that FLT3-ITD and p-STAT5 downregulation in hypoxia is abrogated in the absence of glutamine. **C.** FLT3-ITD AML cells were treated with CHX to block new protein synthesis, with or without addition of the proteasome inhibitor MG-132, and cultured in hypoxia in the presence or absence of 2 mM glutamine. Increased FLT3-ITD proteasomal degradation of occurred in hypoxia in the presence, but not absence, of glutamine.

To test whether restored FLT3 inhibitor cytotoxicity was associated with abrogation of FLT3-ITD and p-STAT5 downregulation, FLT3-ITD AML patient blasts were cultured in hypoxia for 96 hours in medium with and without 2 mM L-glutamine, and expression of FLT3-ITD, p-STAT5 and total STAT was measured by immunoblotting. While expression of FLT3-ITD and p-STAT5 was consistently reduced in cells cultured in glutamine-replete conditions, FLT3-ITD and p-STAT5 expression was preserved, without decrease from Time 0, in cells cultured in hypoxia without glutamine (Figure 5B).

To determine the mechanism by which FLT3-ITD downregulation in hypoxia is abrogated in the absence of glutamine, FLT3-ITD AML patient blasts were treated with CHX to inhibit new protein synthesis, with and without addition of the proteasome inhibitor MG-132, in hypoxia in the presence and absence of 2 mM L-glutamine. FLT3-ITD protein turnover in hypoxia was more rapid in the presence of glutamine than in its absence, and turnover was abrogated by co-culture with MG-132 (Figure 5C), consistent with FLT3 downregulation due to accelerated proteasomal degradation in the presence of glutamine, but not in its absence.

### Glutamine availability regulates c-CBL expression in hypoxia

c-CBL protein expression was measured in FLT3-ITD AML patient blasts cultured in hypoxia for 96 hours in medium with and without 2 mM L-glutamine. c-CBL protein expression was increased in cells cultured with 2 mM L-glutamine, but not in medium without glutamine supplementation (Figure 6). These data suggest that lack of FLT3-ITD downregulation in hypoxia in the absence of glutamine may be due to lack of increase in c-CBL expression, supporting a potential role for c-CBL in facilitating FLT3-ITD proteasomal degradation in hypoxia in the presence of glutamine.

**Figure 6.**
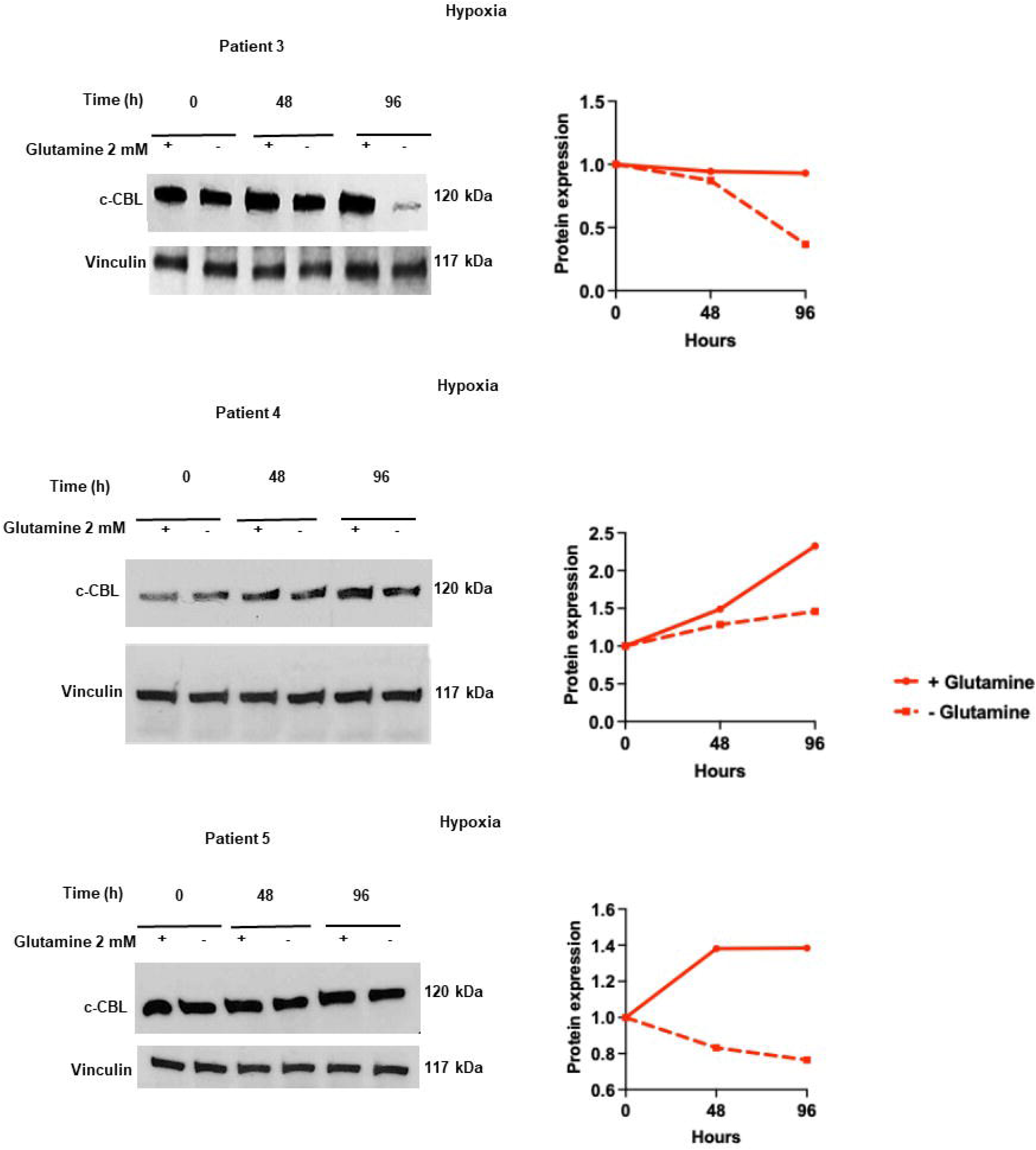
Glutamine deprivation abrogates increased expression of c-CBL in hypoxia. FLT3-ITD AML blasts were cultured in hypoxia for 96 hours in medium with or without 2 mM L-glutamine and immunoblotted for expression of c-CBL and vinculin loading control. Immunoblot and graphic representation are shown. c-CBL protein expression was increased in cells cultured with, but not without, glutamine supplementation.

### The glutaminase inhibitor telaglenastat (CB-839) abrogates FLT3-ITD degradation in hypoxia

Telaglenastat (CB-839) was previously shown to sensitize FLT3-ITD AML cells to quizartinib *in vitro* and *in vivo* (25). To test whether pharmacologic glutaminase inhibition could phenocopy glutamine deprivation, AML cells were treated with telaglenastat (100 nM) for 96 hours in hypoxia. Telaglenastat treatment abrogated c-CBL upregulation and preserved FLT3-ITD protein expression and p-STAT5 signaling in FLT3-ITD AML cell lines and patient samples, abrogating the glutamine-dependent downregulation observed in hypoxia (Figure 7A).

**Figure 7.**
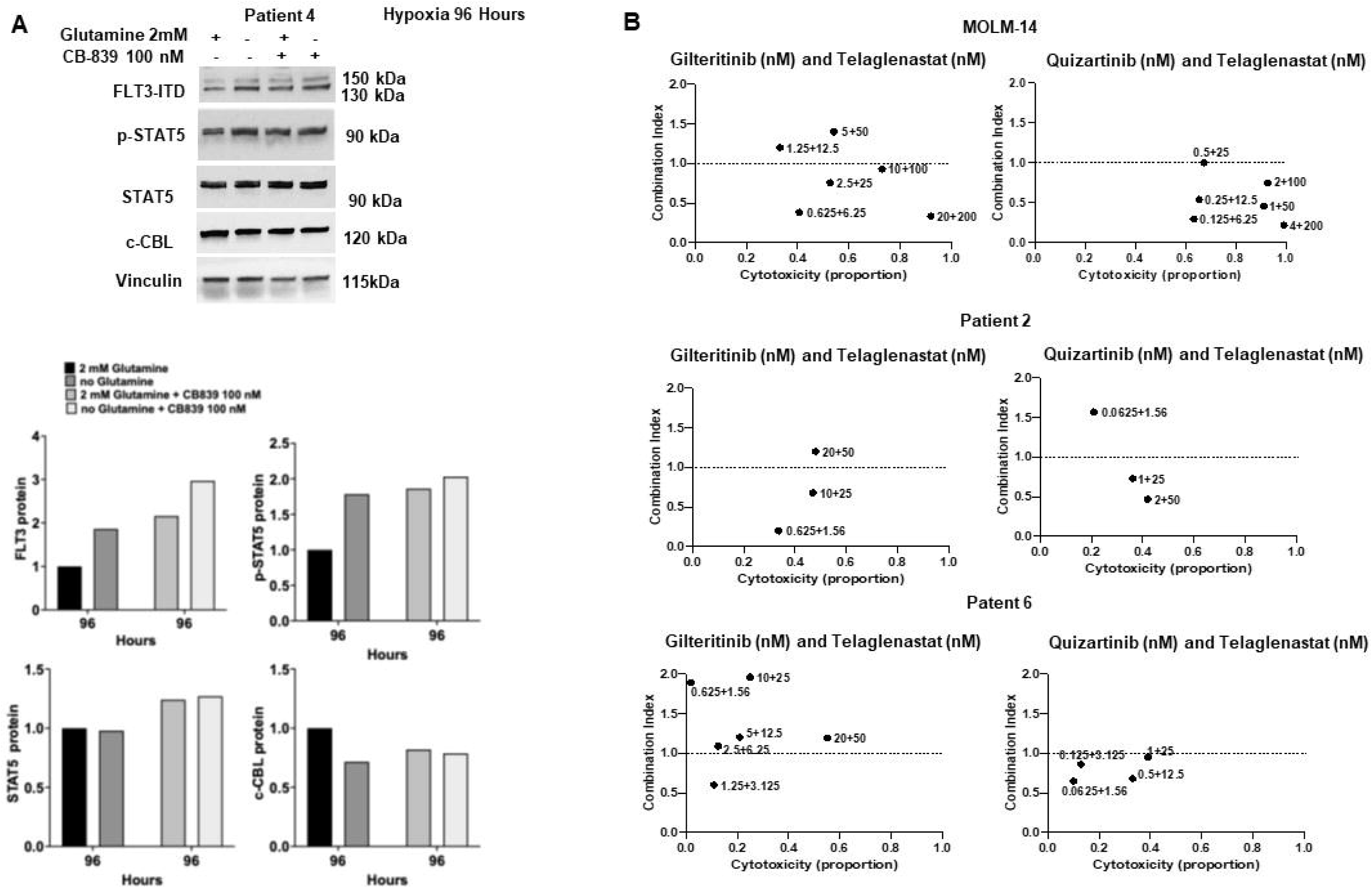
The glutaminase inhibitor telaglenastat (CB-839) abrogates c-CBL upregulation and FLT3-ITD and p-STAT5 downregulation in hypoxia and synergizes with FLT3 inhibitors in FLT3-ITD AML cells. **A.** Primary FLT3-ITD AML cells cultured in hypoxia with and without glutamine (2 mM), with telaglenastat (100 nM) and with glutamine and telaglenastat and harvested at 0, 48 and 96 hours were immunoblotted for c-CBL, FLT3, p-STAT5, STAT5 and vinculin expression, and bands were quantified by densitometry. Immunoblots and densitometric analysis are shown. **B.** MOLM-14 cells and primary FLT3-ITD AML cells from two patients were cultured in hypoxia in the presence of glutamine with telaglenastat and the FLT3 inhibitors gilteritinib or quizartinib at diverse concentrations. Drug combination effects were analyzed using the Chou-Talalay method. Combination index <1, =1 or >1 indicated synergy, additivity and antagonism, respectively. Synergistic effects were seen.

### Telaglenastat synergizes with FLT3 inhibitors in hypoxia

We next assessed whether combining telaglenastat with FLT3 inhibitors produced synergistic cytotoxicity in hypoxia. MOLM-14 cells and FLT3-ITD AML blasts from two patients were cultured in hypoxia with gilteritinib or quizartinib and telaglenastat at diverse concentrations for 48 hours. Synergy of telaglenastat with both FLT3 inhibitors was demonstrated (Figure 7B).

## Discussion

FLT3-ITD AML patient outcomes have improved with incorporation of FLT3 inhibitors into treatment (5), but they remain suboptimal. Notably, treatment with FLT3 inhibitors clears FLT3-ITD AML blasts from the PB, but not the BM (9), a protective niche that impairs FLT3 inhibitor response. FLT3-ITD AML cells persist in the hypoxic BM niche (13–15), and can then also acquire mutations conferring intrinsic FLT3 inhibitor resistance (10). In this study, we demonstrate that FLT3-ITD AML cells acquire resistance to FLT3 inhibitors in hypoxia. Resistance is associated with post-translational downregulation of FLT3-ITD caused by accelerated FLT3-ITD proteasomal degradation, associated with increased expression of the c-CBL E3 ubiquitin ligase (22). We also found that glutamine availability is essential for development of FLT3 inhibitor resistance in hypoxia, as glutamine deprivation or pharmacologic inhibition of glutaminase with telaglenastat (CB-839) prevented c-CBL increased expression, FLT3-ITD downregulation and FLT3 inhibitor resistance in hypoxia. Moreover, telaglenastat synergized with FLT3 inhibitors in hypoxia, supporting future clinical testing.

The E3 ubiquitin ligase c-CBL is a well-established negative regulator of activated receptor tyrosine kinases, including FLT3, targeting them for ubiquitination and proteasomal degradation (22). Missense mutations in c-CBL occur as rare, but recurring, molecular abnormalities in AML and drive leukemia through overexpression of WT FLT3 (26–29). c-CBL mutations may co-occur with FLT3-ITD in AML, and in fact co-occur in the MOLM-13 and MOLM-14, but not MV4-11, FLT3-ITD AML cell lines (30,31). Mice transplanted with hematopoietic stem cells expressing both c-CBL mutations and FLT3-ITD develop aggressive myeloid leukemia, although, surprisingly, FLT3-ITD expression and STAT5 activation are not enhanced in the presence of the c-CBL mutations (32). Of note, c-CBL mutations were not present in any of the AML patient samples that we studied here (Supplementary Table S1), and, in contrast to the mutation-driven models described above, we report upregulation of WT c-CBL expression via a non-mutational, metabolically regulated mechanism in hypoxia, with consequent accelerated FLT3-ITD turnover, downregulation of FLT3-ITD expression and resistance to FLT3 inhibitors. Our findings are also consistent with prior reports that c-CBL regulates FLT3-ITD (22,26), as well as WT FLT3.

Our findings raise an apparent paradox with regard to FLT3-ITD oncogene addiction. FLT3-ITD AML cells are dependent on constitutive FLT3 signaling for survival (4), yet hypoxia-induced FLT3-ITD and p-STAT5 downregulation does not cause apoptosis. This suggests that hypoxia promotes a reversible adaptive state, likely characterized by quiescence, in which FLT3-ITD AML cells reduce reliance on FLT3 signaling. Such a state has been described for AML cells residing in hypoxic BM niches, where reduced metabolic and proliferative activity confers drug tolerance (15). Glutamine appears to enable this adaptive transition by facilitating c-CBL-mediated FLT3-ITD downregulation, while glutamine deprivation or glutaminase inhibition prevents this adaptation, maintaining FLT3-ITD expression and susceptibility to FLT3 inhibition.

Our data expand previous observations on FLT3 downregulation in AML cells in hypoxia. FLT3 expression was previously reported to be variably reduced in AML cells cultured for 48 hours in hypoxia, but not normoxia, in a proteasome-dependent manner, independent of mutational status (33). We observed FLT3 downregulation in hypoxia in all FLT3-ITD samples studied, with downregulation more pronounced after 96 hours than after 48 hours, possibly explaining the more uniform results in our work. In contrast, changes in FLT3 expression were variable among three WT FLT3 AML samples studied here, and were less pronounced, consistent with the prior results (33). We have demonstrated here that hypoxia-induced FLT3-ITD downregulation is not a passive consequence of reduced oxygen tension, but rather an active, metabolically regulated process in AML cells with FLT3-ITD. Specifically, we show that glutamine availability governs this effect through increased expression of c-CBL and consequent ubiquitination and proteasomal degradation of FLT3-ITD.

Mechanistically, increased expression of c-CBL in hypoxia was not associated with upregulation of c-CBL mRNA expression nor altered protein turnover or proteasomal degradation. It is therefore likely mediated by post-transcriptional mechanisms, potentially involving mTOR signaling or hypoxia-regulated microRNAs (34–37). Glutamine regulates mTORC1, suggesting metabolic control of c-CBL accumulation in hypoxia.

While these data support a model in which glutamine availability promotes c-CBL expression through post-transcriptional mechanisms, we cannot exclude the possibility that glutamine deprivation in hypoxia imposes broader metabolic constraints that globally impair protein synthesis. c-CBL regulation might reflect both specific nutrient-sensitive signaling pathways and a more general requirement for bioenergetic support of adaptive protein translation under hypoxic stress.

Functionally, c-CBL is required for FLT3-ITD downregulation in hypoxia, as its silencing restores FLT3 expression, STAT5 signaling, and FLT3 inhibitor sensitivity, identifying it as a key mediator of hypoxia-driven therapeutic resistance. Other CBL family members, including CBL-b, may also contribute to FLT3-ITD degradation (22).

We identify a novel role for glutamine in FLT3-ITD AML. As a key metabolic substrate supporting bioenergetics and redox balance, glutamine sustains survival of FLT3-ITD AML cells under FLT3 inhibition (17,18,38). FLT3 inhibitors impair glucose utilization, rendering tricarboxylic acid (TCA) cycle activity glutamine-dependent, such that co-targeting glutamine metabolism enhances cell death (17). L-asparaginase, which hydrolyzes glutamine, was shown to decrease intracellular glutamine and glutamine-derived TCA metabolites in FLT3-ITD AML cells, potentiate cell death induced by quizartinib, and synergize with quizartinib (18). Finally, gilteritinib additionally downregulated the glutamine transporter SNAT1 (SLC38A1) and decreased glutamine metabolism through the TCA cycle, and CB-839 synergized with gilteritinib (26). We show here that glutamine not only sustains energy metabolism in FLT3-ITD AML cells treated with FLT3 inhibitors, but also enables post-translational FLT3-ITD downregulation in hypoxia, with consequent FLT3 inhibitor resistance.

Notably, the coupling between metabolism and oncogenic protein stability observed here parallels findings in another tyrosine kinase-driven myeloid malignancy. In chronic myeloid leukemia, glutamine availability under hypoxia regulates BCR-ABL protein expression independently of transcriptional control, influencing sensitivity to BCR::ABL inhibitors (39). BCR::ABL is also a c-CBL substrate (40). These observations suggest a broader principle whereby nutrient availability within hypoxic stem cell niches governs oncogenic signaling through post-translational mechanisms, thereby shaping therapeutic response.

These findings highlight the importance of modeling AML under physiologically relevant microenvironmental conditions. Conventional normoxic culture systems fail to capture hypoxia-driven metabolic regulation of oncogenic signaling and may obscure clinically relevant resistance mechanisms. Incorporating hypoxia and nutrient modulation reveals adaptive pathways likely operative *in vivo.* Other approaches to sensitizing FLT3-ITD AML in the BM niche to FLT3 inhibitors have included P13K (41) and AXL (42) inhibition.

Our results provide a strong rationale for therapeutic co-targeting of glutamine metabolism and FLT3 signaling. Glutaminase inhibition was demonstrated to enhance FLT3 inhibitor efficacy through disruption of metabolic homeostasis and induction of oxidative stress (25), and *in vivo* studies (17,25) support the feasibility of this approach. We contribute an additional mechanism by demonstrating that glutaminase inhibition stabilizes FLT3-ITD protein under hypoxia, thereby enhancing target engagement and restoring FLT3 inhibitor sensitivity within the BM niche.

Our results also further support potential therapeutic efficacy of telaglenastat in FLT3-ITD AML. Telaglenastat was previously shown to sensitize FLT3-ITD AML cells to quizartinib *in vitro* and *in vivo* (17,25). Here we further show that telaglenastat restores FLT3 inhibitor sensitivity and synergizes with FLT3 inhibitors in hypoxia. Mechanistically, this interaction is likely largely driven by preserved FLT3-ITD protein expression, thereby sensitizing FLT3-ITD AML cells that persist as MRD in the BM niche to FLT3 inhibitors as an approach to eliminating MRD, rather than by induction of an independent cytotoxic pathway.

Telaglenastat has been well tolerated in clinical trials (43–45). Notably, telaglenastat was combined with azacitidine as a combined metabolic and epigenetic approach in a recent clinical trial in patients with advanced myelodysplastic syndrome, with good tolerability and favorable response rate (44). Our data support future clinical testing in FLT3-ITD AML.

## Acknowledgements

The authors gratefully acknowledge the patients who donated samples under a University of Maryland School of Medicine IRB-approved tissue procurement protocol, University of Maryland Greenebaum Comprehensive Cancer Center clinical research staff for patient sample procurement, the University of Maryland Greenebaum Comprehensive Cancer Genomics Shared Resource for molecular analyses, and Pathology and Biorepository Shared Resource and Translational Laboratory Shared Resource for tissue banking.

## Author Contributions

G.S. and M.R.B. conceptualized the study and designed the experiments. G.S. and AC performed and analyzed the experiments. B.R. contributed to hypoxia studies and provided technical support with the hypoxia system. E.B. provided intellectual input, and supported the study through access to the hypoxia system in his laboratory. G.S. and M.R.B. wrote the manuscript. All authors reviewed, edited, and approved the final version of the manuscript.

## Data Availability

All relevant data are included in the article and the supplementary file. Data are available from the corresponding author upon reasonable request.

