## Supplemental Tables 1 and 2 for "Glutamine-Dependent Downregulation of FLT3-ITD is a Mechanism of FLT3 Inhibitor Resistance in FLT3-ITD AML in Hypoxia"

**Supplemental Material**

**Supplemental Table S1 – Patient Samples**

| **Patient** | **Age, sex** | **Disease status** | **WBC**  **(x10^9^/L)** | **Blasts**  **(%)** | **Karyotype** | ***FLT3*-ITD**  **size**  **(bp)** | ***FLT3*-ITD allelic burden (%)** | **Other mutated genes** |
| --- | --- | --- | --- | --- | --- | --- | --- | --- |
| 1 | 78M | Diagnosis | 36.0 | 60 | 46,XY[20] | 350  366  373 | 9  23  10 | *SF3B1*  *RUNX1* |
| 2 | 60M | Diagnosis | 44.4 | 79 | 46,XY[20] | 355 | 60 | *NPM1,*  *IDH2*  *SRSF2*  *CEPB1* |
| 3 | 60M | Relapse | 45.8 | 41 | 46,XY[6] | 352  361 | 31  3 | *WT1*  *ASXL1*  *RUNX1*  *TET2* |
| 4 | 50F | Diagnosis | 44.3 | 58 | 46,XX[20] | 396 | 87 | *DEK-NUP214* |
| 5 | 69F | Diagnosis | 35.2 | 42 | 46,XX,t(6;9)(p22;q34)[20] | 387 | 55 | *WT1*  *DEK-NUP214* |
| 6 | 60M | Diagnosis | 131.0 | 78 | 46,XY[20] | 356  376 | 84  51 | *DNMT2A*  *TET2*  *NPM1* |
| 7 | 72M | Diagnosis | 41.6 | 68 | 46,XY[20] | - | - | *NRAS*  *NPM1,*  *PTPN11*  *IDH2* |
| 8 | 27F | Diagnosis | 70.3 | 71 | 45,XX,t(6;11)(q27;q23),-7[20] | - | - | *NRAS*  *KRAS* |
| 9 | 71F | Diagnosis | 98.7 | 83 | 46,XX[20] | - | - | *TET2*  *TET2*  *NPM1,*  *SRSF2* |

**Supplemental Table S2 - Primers**

| FLT3 forward | 5^I^ TGTCGAGCAGTACTCTAAACA 3^I^ |
| --- | --- |
| FLT3 reverse | 5^I^ ATCCTAGTACCTTCCCAAACTC 3^I^ |
| c-CBL forward | 5^I^ TCAGCCTAGGCGAAACCTAA 3^I^ |
| c-CBL reverse | 5^I^ TCCCTCTAGGATCAAACGGA 3^I^ |
| GAPDHF forward | 5^I^ ACCATGGAGAAGGCTGGGGCTCAT 3^I^ |
| GAPDHF reverse | 5^I^ ACAGTTTCCCGGAGGGGCCAT 3^I^ |
